# The genome sequence of the reef-building coral *Cyphastrea microphthalma* from the Persian/Arabian Gulf

**DOI:** 10.64898/2026.11.26.769504

**Authors:** Rachel Alderdice, Abdoallah Sharaf, Anna Fiesinger, Gabriela Perna, Hannah Manns, John A. Burt, Christian R. Voolstra

**Author notes:** **Corresponding authors:** Rachel Alderdice, Christian R Voolstra.

## Abstract

We present a genome assembly from the coral species *Cyphastrea microphthalma* from the southern Persian/Arabian Gulf (Cnidaria; Anthozoa; Hexacorallia; Scleractinia; Merulinidae). The assembly is 681.2 Mb in size, spanning 5,554 scaffolds with a contig N50 of 244.4 kb. The assembly has a BUSCO completeness of 94.1% (single = 85.1%, duplicated = 9%, fragmented = 1.6%, missing = 4.3%) using the metazoa_odb10 reference set (n = 954). Genome annotation identified 24,716 protein-coding genes (including 10 pseudogenes) and a high repeat content of 63.4%. The mitochondrial genome has also been assembled and is 17.9 kb in length.

## Introduction

*Cyphastrea microphthalma*, (Lamarck, 1816) (Eukaryota; Metazoa; Eumetazoa; Cnidaria; Anthozoa; Hexacorallia; Scleractinia; Merulinidae; NCBI:txid570133) is a common massive or encrusting reef building scleractinian coral in the family Merulinidae (Fig. 1). The species is distributed throughout the tropical Indo Pacific, including the Red Sea, Persian Gulf, Gulf of Oman, Southeast Asia, Australia, Japan, and West/Central Pacific [1–5]. Colonies typically form irregular mounds but adopt a more encrusting growth form in low-light environments, such as turbid waters. These coral tend to occupy reef flats, channels, lagoon, and forereef slopes at depths of 6-25 m. Key diagnostic skeletal features include tall, conical corallites with primary corallites possessing 10 primary septa. Brown, cream to green visual color morphs, often with white septa, have been observed [6]. Like most corals, this species is a simultaneous hermaphrodite that broadcast-spawns annually over one or few consecutive nights [7,8].

**Figure 1.**
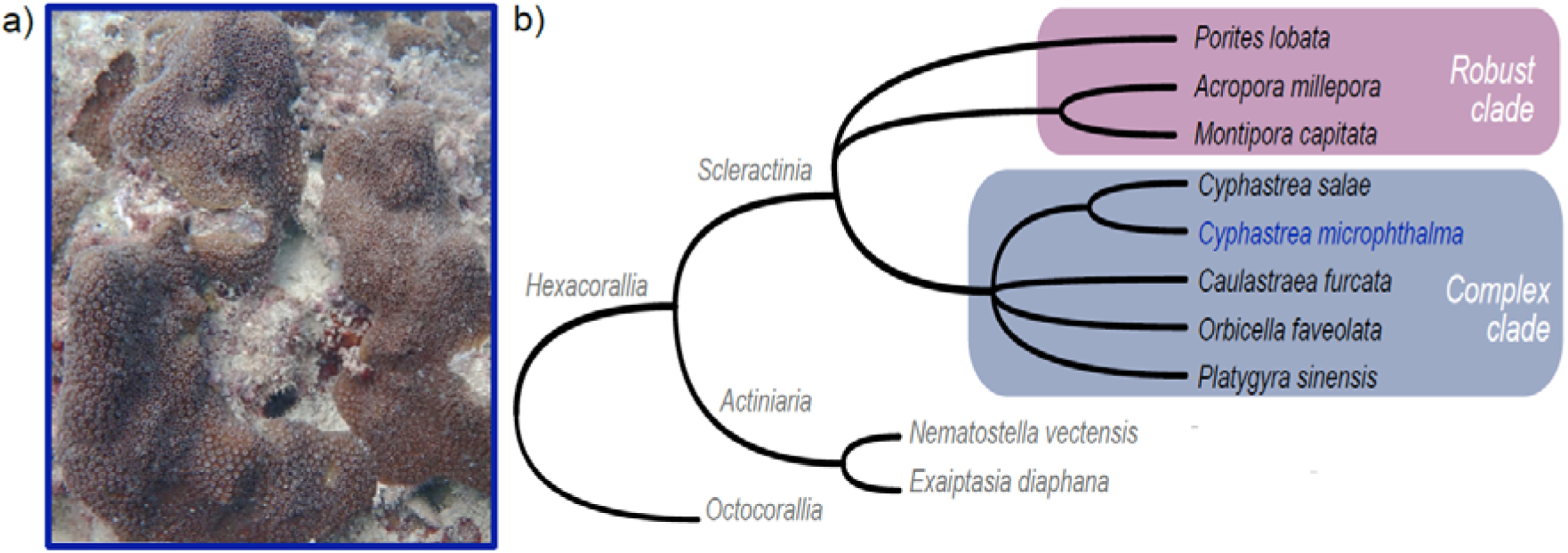
The species *Cyphastrea microphthalma* (Lamarck, 1816). **(a)** *In situ* colony photo of *Cyphastrea microphthalma* from Saada Reef in the Persian/Arabian Gulf. Photo credit: Christian R. Voolstra. **(b)** Cladogram depicting the evolutionary relationship of *Cyphastrea microphthalma* in reference to other Hexacorallia with sequenced genomes.

This species has been reported as a relatively stress-tolerant coral, able to persist in extreme environments such as the world’s warmest ocean body in the Persian/Arabian Gulf or highly fluctuating conditions (turbidity, temperature, macrotides) found in Western Australia [1,9]. Such corals have managed to withstand successive bleaching events, unlike many other species [1,2,9] and can harbor algal endosymbionts such as *Cladocopium thermophilum* and *Durusdium* sp. that confer resilience to thermal stress and high salinity in marginal reef habitats [10].

Corals of the genus *Cyphastrea* are reported to belong to the robust clade (Fig. 1) of the complex-robust deep evolutionary split within hexacorallia [11], based on mitochondrial cytochrome c oxidase subunit I and nuclear ribosomal DNA markers [12]. From this genus, only the species of *Cyphrastea salae* has a scaffold and chromosome level genome publically available on NCBI (https://www.ncbi.nlm.nih.gov/datasets/genome/GCA_964194085.1/; [13]). Given that genomic data, including gene inventory, regulatory features, and structural variants, can provide important insight into coral adaptation to climate change [14], and that genome size and architecture vary substantially across coral taxa [15], expanding genomic resources to underrepresented lineages from extreme environments is important for enabling more comprehensive evolutionary and functional comparisons.

## Results and Discussion

### Genome Assembly and Repeat Elements Identification

The genome was sequenced from a single *Cyphastrea microphthalma* colony collected from Al Saada Reef (SA) in the southern Persian/Arabian Gulf on 1^st^ June 2022 (Table 1). Following a hybrid approach, ONT long-reads were used for the initial assembly then short paired-end reads were used for polishing. A total of 17,396,181 ONT long-reads were generated from an R10.4.1 platform (Oxford Nanopore Technologies, Oxford, UK). After adapter removal and trimming, the mean and median read lengths were 5,933 and 3,246 bp, respectively, with a read-length N50 of 12,281 bp. The reads showed a mean basecall quality score of 14.4, and the longest read sequenced was 40 kbp with a quality score of 19.6. Mean sequencing coverage was estimated at ∼136.8X for the ONT long reads and ∼21.2X for the short reads. The estimated genome size was determined using a k-mer approach (k = 31) with the short reads, resulting in an estimated genome size of 590 Mb. The diploid genome assembly is predominantly homozygous (99.1% AA) and exhibits minimal heterozygosity (0.86% AB) (Fig. 2). Although Smudgeplot v0.4.0 strongly supported a diploid genome, our short-read sequencing data were insufficient for reliable ploidy inference from the GenomeScope v2.0 k-mer distribution (Fig. 2). The genome size estimated in this study aligns with other *Cyphastrea* genome sizes inferred using the Genomes on a Tree (GoaT) platform (https://goat.genomehubs.org).

**Figure 2.**
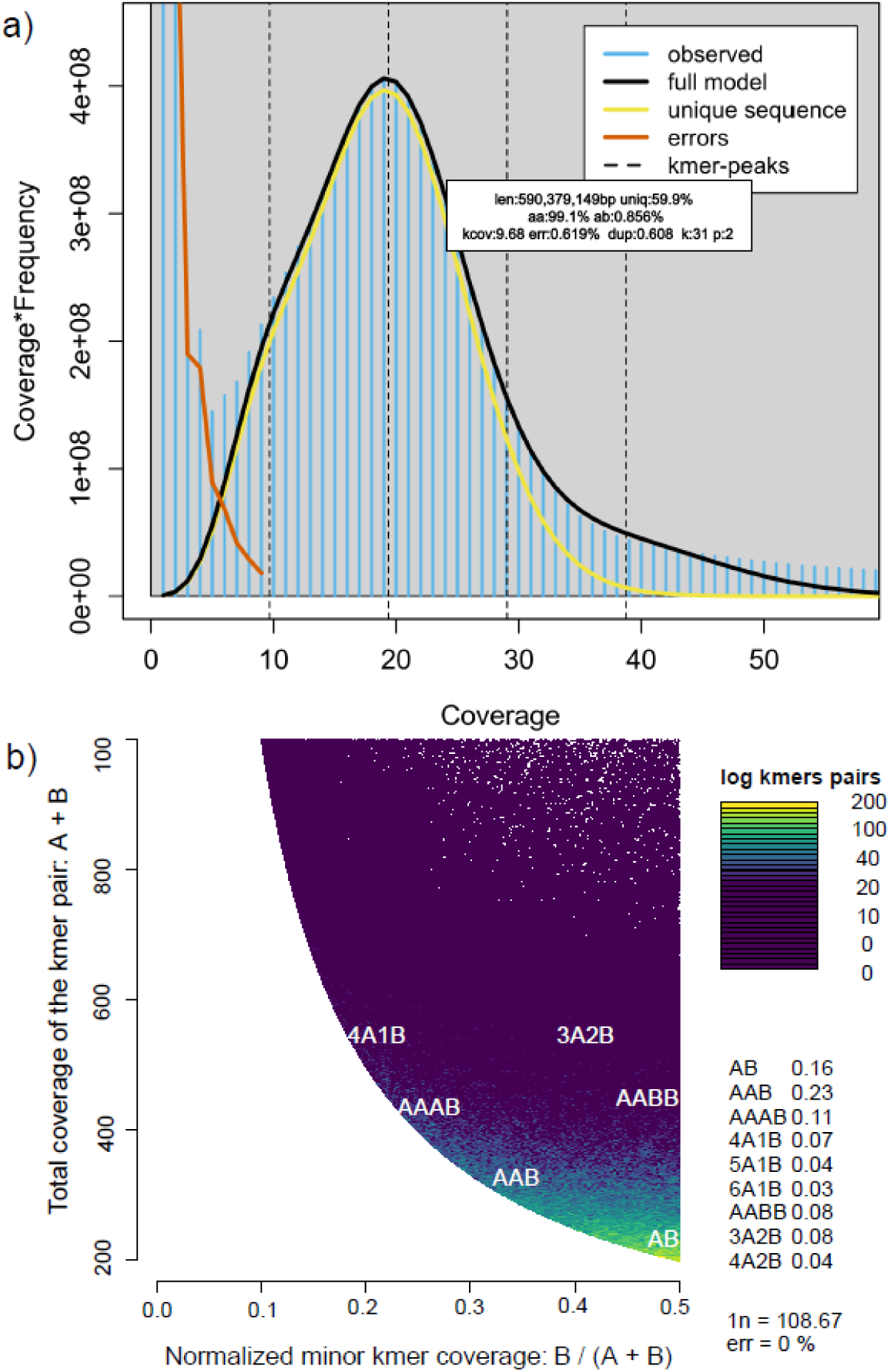
K-mer profiling of the *Cyphastrea microphthalma* genome based on the Illumina short-read dataset. **(a)** The GenomeScope v2.0 analysis shows the genome size estimation to be 590 Mb, with high homozygosity (AA 99.1%; AB 0.85%). The following abbreviations are used in the diagram: len – inferred total genome length, uniq – percent of the genome that is unique (not repetitive; 59.9%), AB – overall rate of heterozygosity, kcov – mean k-mer coverage for heterozygous bases (9.68), err – error rate of the reads (0.62%), dup – average rate of read duplications (0.61), k – k-mer size (31). **(b)** The Smudgeplot v0.4.0 analysis shows the distribution of k-mer pair multiplicities on a log10 scale. The Smudgeplot is a visual representation of the diploidy level that allows for the visualization of heterozygosity.

**Table 1.**
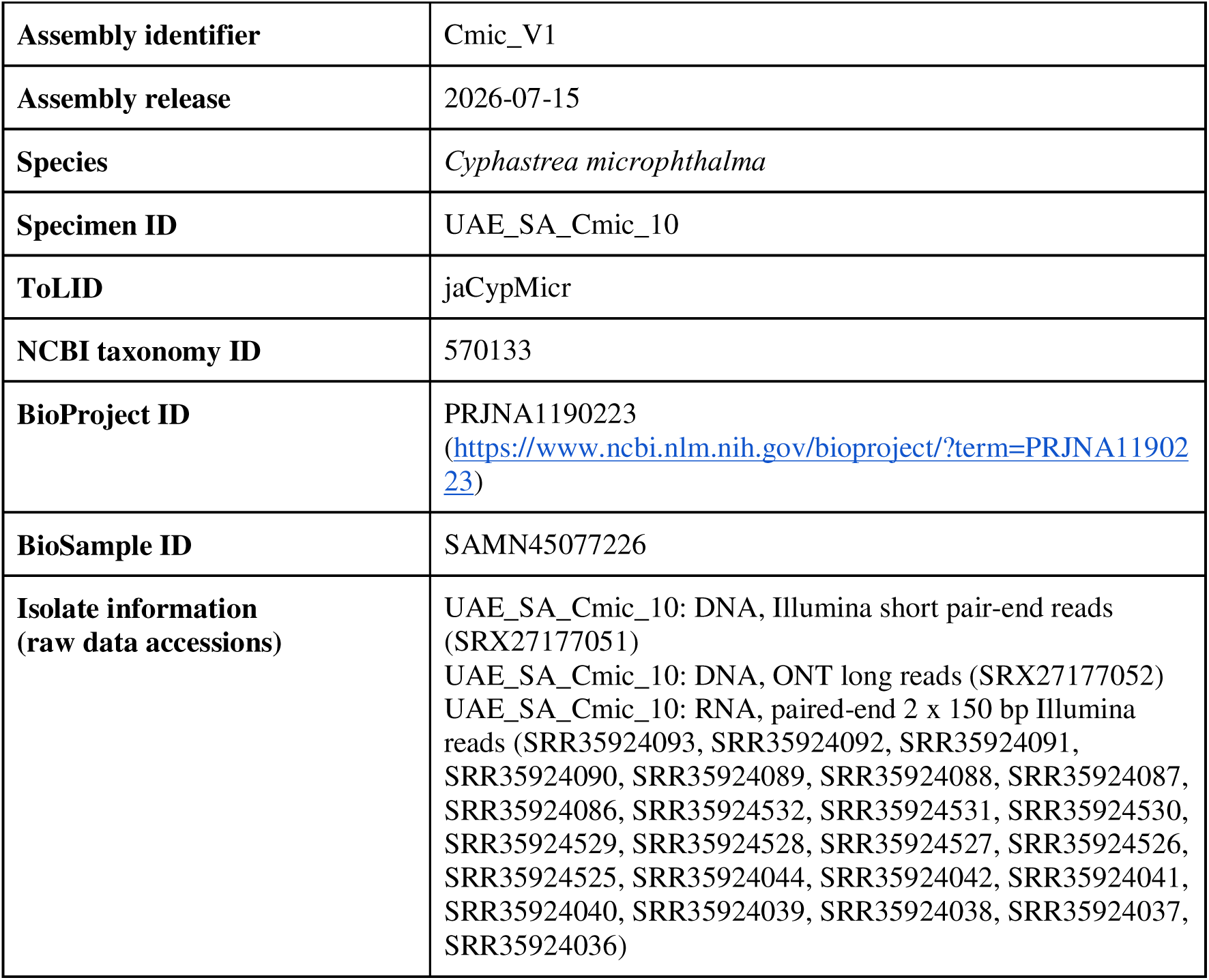
Project accession data for *Cyphastrea microphthalma*.

| **Assembly identifier** | Cmic_V1 |
|---|---|
| **Assembly release** | 2026-07-15 |
| **Species** | *Cyphastrea microphthalma* |
| **Specimen ID** | UAE_SA_Cmic_10 |
| **ToLID** | jaCypMicr |
| **NCBI taxonomy ID** | 570133 |
| **BioProject ID** | PRJNA1190223 (https://www.ncbi.nlm.nih.gov/bioproject/?term=PRJNA1190223) |
| **BioSample ID** | SAMN45077226 |
| **Isolate information<br>(raw data accessions)** | UAE_SA_Cmic_10: DNA, Illumina short pair-end reads (SRX27177051)<br>UAE_SA_Cmic_10: DNA, ONT long reads (SRX27177052)<br>UAE_SA_Cmic_10: RNA, paired-end 2 x 150 bp Illumina reads (SRR35924093, SRR35924092, SRR35924091, SRR35924090, SRR35924089, SRR35924088, SRR35924087, SRR35924086, SRR35924532, SRR35924531, SRR35924530, SRR35924529, SRR35924528, SRR35924527, SRR35924526, SRR35924525, SRR35924044, SRR35924042, SRR35924041, SRR35924040, SRR35924039, SRR35924038, SRR35924037, SRR35924036) |

The initial draft assembly spanned 780.2 Mb. Following assembly quality assessment and contamination screening, 285 contigs identified as potential contaminants based on their GC content, sequencing coverage, and taxonomic assignment were removed. The resulting filtered assembly spanned 681.2 Mb across 5,827 contigs, with a contig N50 of 244.4 kb and a GC content of 39.5% (Table 2). Taxonomic assessment of the filtered assembly showed that the retained contigs were consistent with the expected taxonomic affiliation of the target organism (Cnidaria), with no detectable evidence of substantial contamination from non-target taxa (Fig. 3). The genome size is similar to that of *C. salae* (656 Mb; <u>GCA_964194085.1</u>). Following scaffolding, the assembly comprised 5,554 scaffolds with a scaffold N50 of 266.7 kb and a maximum scaffold length of 2.8 Mb (Fig. 3). The mitochondrial genome was assembled into a single contig of 17.9 kb, similar to the length of the *C. serailia* mitogenome [16], with 16 protein-coding genes, 4 tRNAs, and 1 rRNA (Fig. 4).

**Figure 3.**
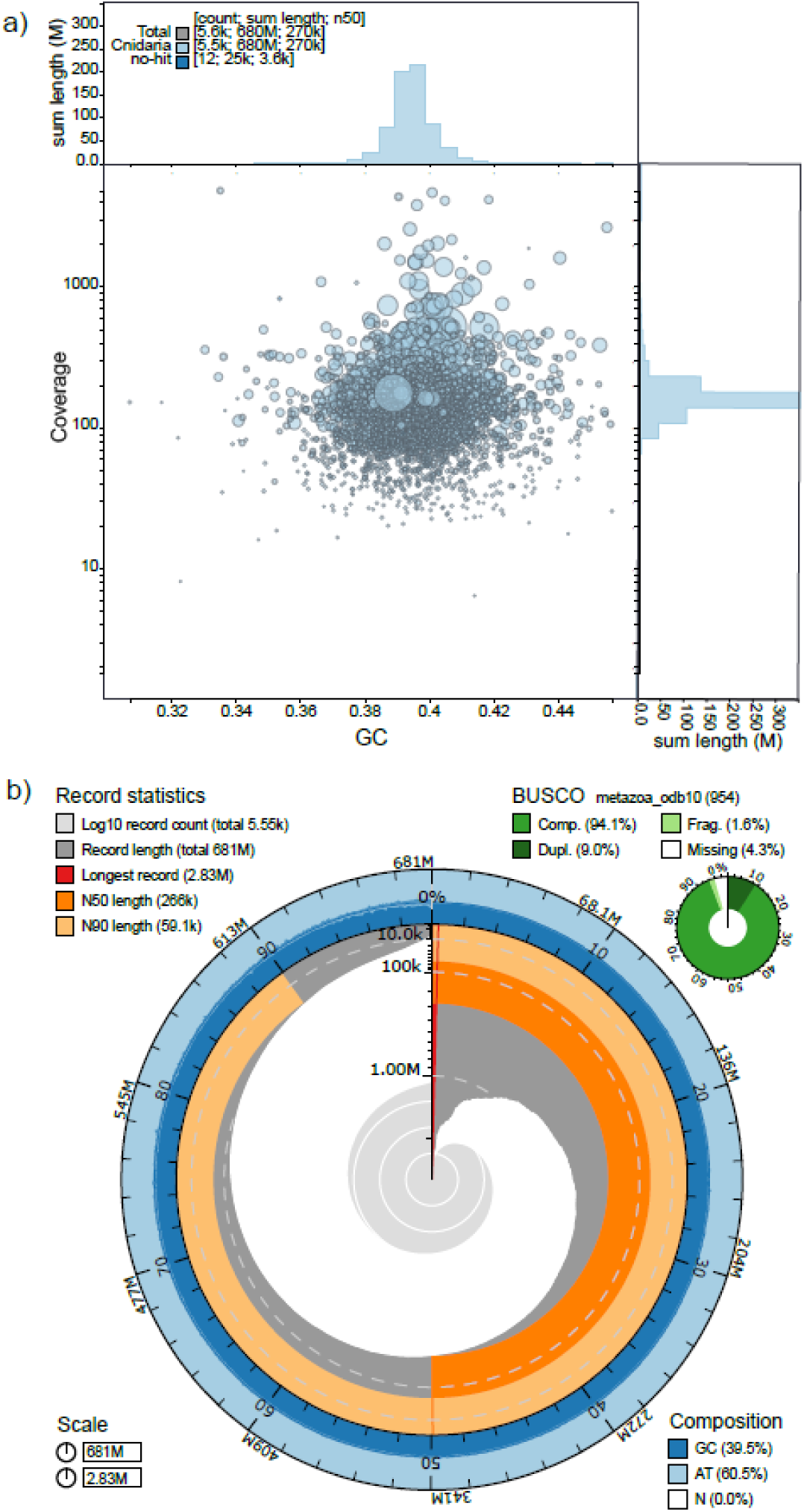
The nuclear genome assembly of *Cyphastrea microphthalma*. **(a)** Circle-binned blob plot of GC proportion showing no evidence of contamination by non-target species after filtering. Coverage is based on long read sequences. Scaffolds are coloured in light blue for the phylum Cnidaria and dark blue for no-hit, and circles are sized in proportion to scaffold length. Histograms show the distribution of scaffold length sums along each axis. **(b)** Key assembly metrics. The BlobToolKit v4.3.0 snail plot comprises 1,000 size-ordered bins arranged around the circumference, with each bin representing 0.1% of the 681,218,052 bp assembly. The distribution of scaffold lengths is shown in dark grey with the plot radius scaled to the longest scaffold present in the assembly (2,834,921 bp, red). Dark orange and light orange arcs indicate N50 (266,700 bp) and N90 (59,100 bp) scaffold sizes, respectively. The central light grey spiral represents the cumulative scaffold count on a logarithmic scale, with white scale lines showing successive orders of magnitude. The dark and light blue outer rings show the composition of GC, AT, and N percentages. A summary of complete, fragmented, duplicated, and missing BUSCO genes, assessed in genome assembly mode metazoa_odb10 dataset (n = 954 BUSCOs), is shown in the top right.

**Figure 4.**
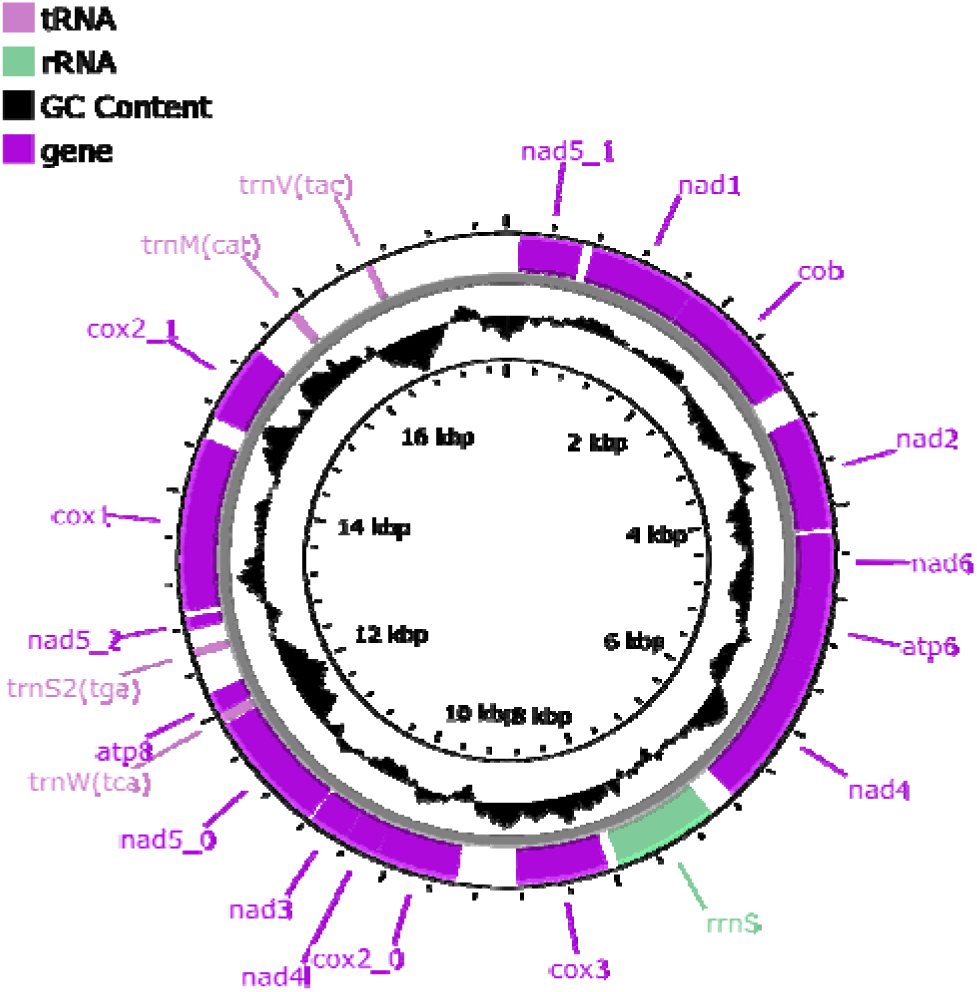
Mitochondrial genome of *Cyphastrea microphthalma*. The annotated features are indicated by color: light violet are the annotated genes (n = 16), light green are rRNAs (n = 1), black are GC content and pink are tRNAs (n = 4). Circularized = TRUE.

**Table 2.** Genome assembly data for *Cyphastrea microphthalma*.

| **Assembly accession number** | GCA_059441185.1<br>(https://www.ncbi.nlm.nih.gov/datasets/genome/GCA_059441185.1/) |
|---|---|
| **Nuclear genome size (Mb)** | 681.2 |
| **Number of contigs** | 5,827 |
| **Contig N50 length (kb)** | 244.4 |
| **Number of scaffolds** | 5,554 |
| **Scaffold (N50) length (kb)** | 266.7 |
| **Longest scaffold (Mb)** | 2.8 |
| **GC content (%)** | 39.5 |
| **Repeat content (%)** | 63.4 |
| **Assembly-level BUSCO<br>(metazoa_odb10, n =954)<br>Completeness (%)<br>-Single-copy (%)<br>-Duplicated (%)<br>Fragmented (%)<br>Missing (%)** | 94.1<br>85.1<br>9.0<br>1.6<br>4.3 |
| **Mitochondrial genome size (kb)** | 17.9 |

### Gene Prediction And Functional Annotation

Structural annotation was done using the GeneForge v1.0 Nextflow pipeline [17], which integrates BRAKER v3.0.8 and funannotate v1.8.5 for eukaryotic gene prediction and automatically selects the highest-quality annotation based on BUSCO v6.0.0 completeness. The BRAKER annotation showed the highest BUSCO completeness and was therefore selected as the final gene set [18,19]. RNA-Seq data from fragments of the same coral colony subjected to different temperature treatments generated a total of 395,989,690 Illumina pair-end reads and were used as transcript evidence for structural annotation within the funannotate pipeline [20]. The final annotation comprised 24,716 genes, including 10 pseudogenes and 1,383 high-confidence tRNA genes. Of which, 21,249 were functionally annotated (Table 3). The number of genes falls within the range of 20,000–40,000 reported for other coral gene sets [14,21,22]. Assembly-level BUSCO analysis indicated a genome completeness of 94.1%, with 1.6% fragmented and 4.3% missing genes based on the metazoa_odb10 reference dataset (*n* = 954). Comparable completeness was obtained using the more taxonomically specific anthozoa_odb12 dataset (*n* = 3,649), with 94.7% (3,455) complete, 2.2% (80) fragmented, and 3.1% (114) missing orthologs. After structural annotation, BUSCO analysis on the predicted gene set revealed 94.4% (901) annotation completeness, 1.9% (18) fragmented, and 3.7% (35) missing genes based on the metazoa_odb10 reference dataset (n = 954). While using anthozoa_odb12, BUSCO results showed 90.8% (3312) complete with 2.6% (95) fragmented and 6.6% (242) missing. The mean gene and protein length was 8,528.6 bp and 456.4 aa, respectively (Table 3). Repetitive sequences comprised a total of 431.50 Mb (63.4%) of the assembly, of which 109.6 Mb (16.1%) were retroelements, 59.5 Mb (8.7%) were DNA transposable elements, and 251.4 Mb (36.9%) were unclassified repeats (Fig. 5). Repeat content was substantially higher than that reported for the *C. salae* genome (40.8%; [13]), suggesting considerable variation in repetitive sequence abundance even among closely related *Cyphrastrea* species. The discrepancy between the k-mer-based genome size estimate and the final assembly span may partly reflect the high repetitive content of the genome, as highly repetitive sequences can contribute to genome size underestimation in k-mer-based approaches [23]. The relatively low short-read coverage may further limit estimation accuracy. Despite the high repeat content (63.4%), the number of protein-coding genes (24,716) remained within the range commonly reported for scleractinian corals (∼20,000-40,000; [14,21,22]) and similar to *C. salae* (27,298), suggesting that the larger repeat content is not accompanied by a concomitant expansion of the protein-coding gene repertoire. More than one third of the assembly (36.9%) consisted of unclassified repeats, highlighting the substantial proportion of repetitive elements that remain poorly characterised and may include lineage-specific repeat families.

**Figure 5.**
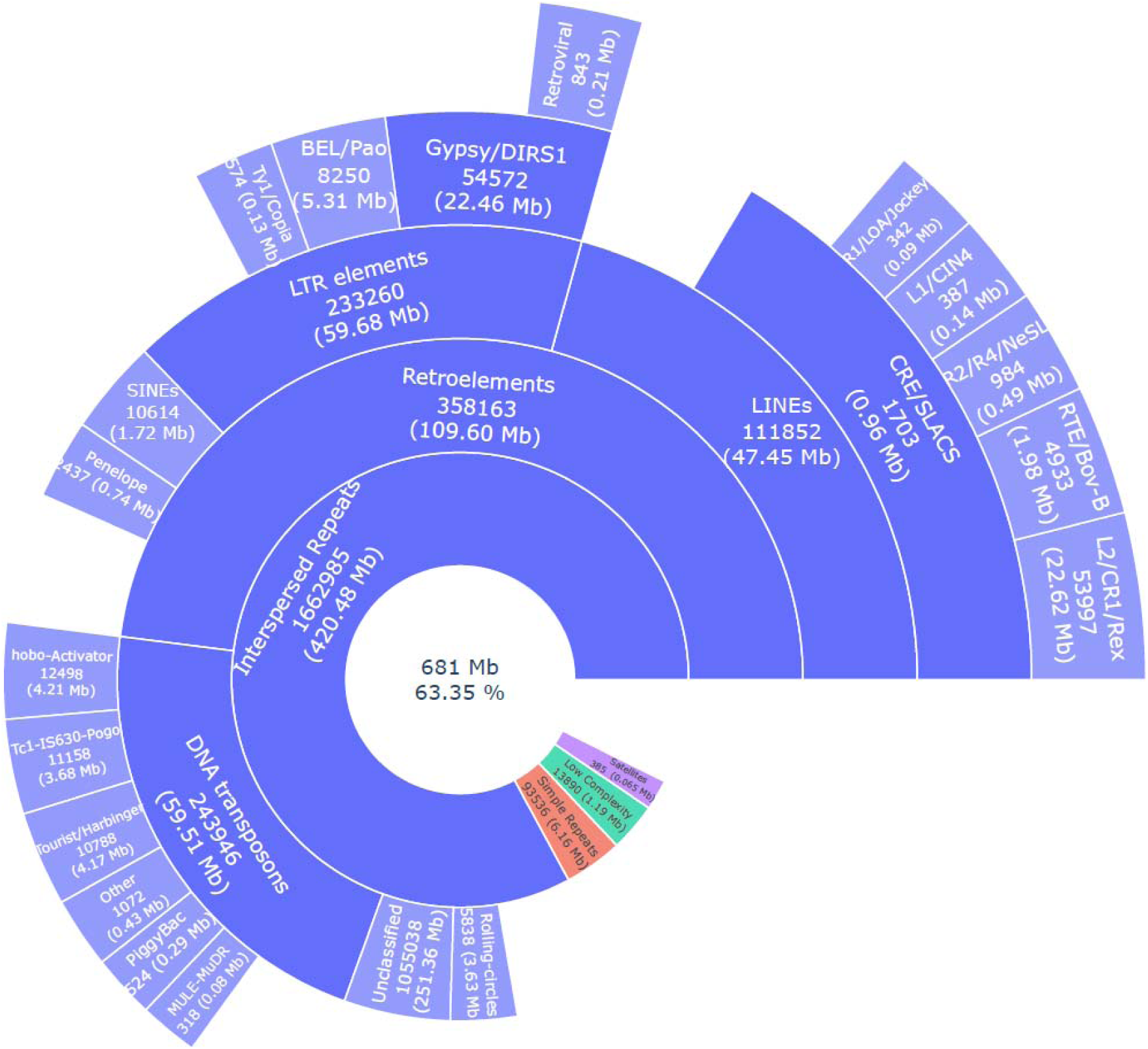
Repeat content of the *Cyphastrea microphthalma* genome. The sunburst plot shows that repetitive sequences account for 431.5 Mb (63.4%) of the 681-Mb genome assembly. Unclassified repeats account for 251.36 Mb (36.9%) of the assembly, whereas DNA transposons account for 59.5 Mb (8.7%).

**Table 3.**
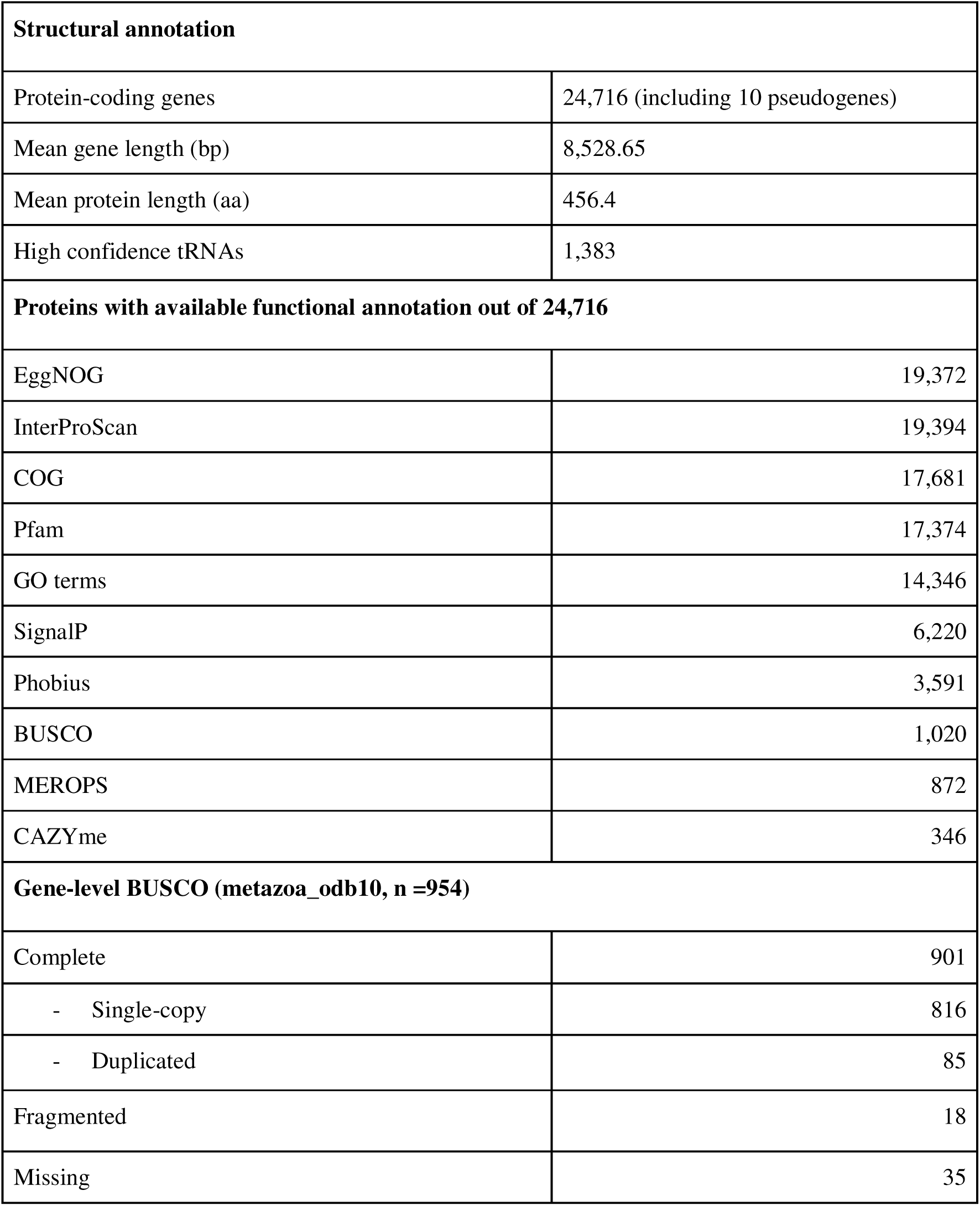
Genome annotation statistics of *Cyphastrea microphthalma*.

| **Structural annotation** |  |
|---|---|
| Protein-coding genes | 24,716 (including 10 pseudogenes) |
| Mean gene length (bp) | 8,528.65 |
| Mean protein length (aa) | 456.4 |
| High confidence tRNAs | 1,383 |
| **Proteins with available functional annotation out of 24,716** |  |
| EggNOG | 19,372 |
| InterProScan | 19,394 |
| COG | 17,681 |
| Pfam | 17,374 |
| GO terms | 14,346 |
| SignalP | 6,220 |
| Phobius | 3,591 |
| BUSCO | 1,020 |
| MEROPS | 872 |
| CAZYme | 346 |
| **Gene-level BUSCO (metazoa_odb10, n =954)** |  |
| Complete | 901 |
| - Single-copy | 816 |
| - Duplicated | 85 |
| Fragmented | 18 |
| Missing | 35 |

## Materials and Methods

### Sample collection

A total of 8 samples (∼2 cm diameter cylinders) were drilled from a single colony (specimen ID: UAE_SA_Cmic_10) of *Cyphastrea microphthalma* at Al Saada Reef (SA; 24.085250, 52.243333) in the southern Persian/Arabian Gulf on 1^st^ June 2022 using a Nemo underwater drill (Nemo Power Tools LLC, Nevada, USA) fitted with a diamond tip corer. Coral fragments were subjected to a short-term acute heat stress assay using the Coral Bleaching Automated Stress System (CBASS; as per [24,25]). Two coral fragments were subjected to one of the four temperature treatments; ‘baseline’ (Maximum monthly mean temperature; MMM, 34.1°C), +4°C, +6°C and +9°C. A fragment per treatment was sampled at two different time points; after the heat hold (360 mins) and at the end of the experiment (1080 mins) after the overnight recovery phase. All fragments were submerged in 3 ml of DNA/RNA Shield (Zymo Research Europe GmbH, Freiburg, Germany) in a Whirlpak bag for at least 3 hours at RT then tissue was airpicked off the skeleton using an air pump with sterile filter tips. Tissue slurry was transferred to 2 ml cryotubes and stored at 4 °C until shipment to the lab facilities at the University of Konstanz, Germany. The samples were stored at -20 °C for subsequent DNA/RNA extractions. All samples were considered for total RNA extractions and the baseline temperature treatment sample were also used for DNA extraction.

### Nucleic acid isolation and sequencing

DNA was extracted using the DNeasy Blood & Tissue kit (Qiagen, Hilden, Germany) according to the manufacturer’s protocol. Briefly, following homogenization, debris was spun down at 5,000 x *g* for 3 min and 90 µl of the supernatant was transferred to a 1.5 ml tube. Subsequently, 90 µl of ATL buffer and 20 µl of Proteinase K (Qiagen, Hilden, Germany) were added and the sample was lysed for 1 hr at 300 rpm. Following the spin column based extraction, two elution steps were applied according to [26] consisting of 2 x 50 µl of AE elution buffer (Qiagen, Hilden, Germany) incubated at 1 and 5 mins respectively, and then centrifuged at 7,000 x *g* for 1 min. The resulting DNA quality and quantity were assessed using a NanoDrop2000 spectrophotometer (Thermo Fisher Scientific, Waltham, Massachusetts, USA). For long-read sequencing, a total of 3.38 µg of DNA was submitted to the NGS Competence Center Tübingen (NCCT) for library preparation using the Oxford Nanopore Technologies (ONT) ligation sequencing kit (SQK-LSK114). Sequencing was performed using an R10.4.1 flow cell (Oxford Nanopore Technologies, Oxford, UK) and base calling was conducted using MinKNOW v23.07.12 (Oxford Nanopore Technologies, Oxford, UK) and Dorado v0.7.4 [27]. For short-read sequencing, DNA was submitted to NCCT for preparation of a PCR-free DNA tagmentation 2 x 150 bp library, followed by whole-genome sequencing on an Illumina NovaSeq 6000 using the S4 Reagent Kit v1.5.

Total RNA was extracted using the RNeasy Mini Kit (Qiagen, Hilden, Germany) adapted for the use with a QIAcube (Qiagen, Hilden, Germany) and low RNA concentrations [28]. The quantity and quality of the RNA was estimated using a NanoDrop2000 spectrophotometer (Thermo Fisher Scientific, Waltham, Massachusetts, USA) as well as gel electrophoresis for quality assessment via the presence of intact 18S and 28S ribosomal RNA bands. The extracted RNA was sent to NCCT for library preparation using the NEBNext Ultra II Directional mRNA kit and subsequent paired-end sequencing on an Illumina NovaSeq (2 x 150 bp).

### Estimation of genome size

The genome size was estimated using a k-mer histogram generated with FastK v1.1 [29] based on the Illumina short reads and visualized in GenomeScope2.0 (RRID:SCR_017014) and Smudgeplot v0.4.0. We note that the K-mer-based estimates used here are not a substitute for experimental validation, which were not possible with these preserved coral tissue samples.

### Nuclear genome assembly, curation, and evaluation

Base-called reads were trimmed using porechop v0.2.4 (RRID:SCR_016967) [30], and the quality was assessed using NanoPlot v1.44.1 (RRID:SCR_024128) [31]. Following a hybrid assembly approach, long reads were used for an initial draft assembly using Flye v2.9.5 (RRID:SCR_017016) [32]. Assembly quality and potential contamination were assessed using BlobToolKit v4.3.0 (RRID:SCR_025882). Taxonomic assignments were supported by nucleotide sequence similarity searches against the NCBI nucleotide (nt) database using BLAST v2.16.0 (RRID:SCR_004870) and translated protein similarity searches using DIAMOND v2.1.10 (RRID:SCR_016071) against the UniProt Reference Proteomes database (RRID:SCR_018666). Contigs were evaluated based on their taxonomic assignments, GC content, and sequencing coverage. Both searches were performed with an E-value threshold of 1 × 10 ². The maximum number of target sequences was set to 10 for BLASTn and 1 for DIAMOND. Sequencing coverage was assessed by mapping the sequencing reads to the initial assembly using Minimap2 v2.28-r1209 (RRID:SCR_018550) [33], followed by BAM sorting with SAMtools v1.21 (RRID:SCR_002105) [34]. The resulting coverage information was incorporated into the BlobToolKit assessment, together with GC content and taxonomic assignments, to identify potential contaminant contigs. Based on the combined evidence from these metrics, 285 contigs identified as potential contaminants and taxonomically inconsistent with the target organism were removed prior to downstream analyses. For polishing, both long and short reads were used with Medaka v1.12.0 and Pilon v1.24 (RRID:SCR_014731), respectively. Duplicated contigs and those shorter than 200 bp were removed using funannotate v1.8.5 (RRID:SCR_023039) [20]. Redundans v2.0.1 was used for final scaffolding. The quality of the final assembly was assessed using BUSCO v6.0.0 (RRID:SCR_015008) [35] and the BlobToolKit v4.3.0 (RRID:SCR_025882).

### Mitochondrial genome assembly and annotation

The mitogenome contig was identified and extracted from the scaffolded and polished assembly graph generated from total DNA sequencing using the de novo assembler Flye v2.9.5 (RRID:SCR_017016), with GetOrganelle v1.7.7.1 (RRID:SCR_022963). The mitogenome was then annotated with MITOS2 v2.1.9 [36]. Mitochondrial genome annotation was performed using MITOS2, an established computational approach for identifying mitochondrial genes. However, repetitive regions and potential gene rearrangements, particularly involving tRNAs, can complicate mitochondrial genome assembly and annotation. Although comparative synteny analyses and experimental validation could provide additional support for mitochondrial genome structure, these were beyond the scope of the present study. The mitogenome is reported as a draft annotation that will be useful for future comparative work.

### Repeat annotation

Genomic DNA repeats were identified using a combination of RepeatModeler v2.0 (RRID:SCR_015027) [37] and the Extensive De novo TE Annotator (EDTA) v2.2.2 pipeline (RRID:SCR_022063) [38], and masked using RepeatMasker v4.1.6 (RRID:SCR_012954) [39]. In addition to the species-specific repeats identified de novo in this genome using RepeatModeler v2.0 and EDTA v2.2.2, repeat elements from closely related Merulinidae genomes (*Orbicella franksi* [GCA_964199315.1], *Cyphastrea salae* [GCA_964194085.1], *Orbicella faveolata* [GCA_002042975.1], *Platygyra sinensis* [GCA_019787425.1], and *Caulastraea furcata* [GCA_965607595.1]) were included to build a broader reference library to identify and mask repeats that may not be detected or fully characterised from the focal genome alone. We acknowledge that localized overmasking cannot be entirely excluded when using repeat libraries from closely related genomes. To visualise the distribution of the identified repeat elements, a sunburst plot was created using Plotly Express library in Python (Plotly Technologies Inc.).

### Gene prediction and annotation

Genome annotation was performed using our fully automated pipeline, GeneForge v1.0 [17]. In brief, both BRAKER v3.0.8 (RRID:SCR_018964) and funannotate v1.8.5 (RRID:SCR_023039) [18,19] were used for structural annotation and gene prediction, generating two sets of predicted gene models and protein sequences. GeneForge evaluated these gene sets using BUSCO completeness. The annotation achieving the highest BUSCO completeness score was selected for functional annotation. GeneForge v1.0 was run on the masked genome assembly, with the RNA-seq alignments and protein evidence serving as complementary inputs. The resulting annotation was subsequently augmented with untranslated regions (UTRs) using the stringtie2utr.py script from BRAKER v3.0.8 (RRID:SCR_018964). Paired-end RNA-seq libraries, generated using a reverse-stranded library preparation protocol, were evaluated for sequence quality using FastQC v0.12.1 (RRID:SCR_014583) and subsequently trimmed to remove adapter sequences and low-quality bases using Trimmomatic v0.39 (RRID:SCR_011848) [40]. Following trimming, 379,942,229 read pairs were retained from 395,989,690 initial read pairs, corresponding to a read-pair retention rate of 95.95%. The quality-filtered reads were mapped to the *C. microphthalma* genome assembly using STAR v2.7.11b (RRID:SCR_004463) [41]. Overall, 60.35% of input reads mapped to the *C. microphthalma* genome assembly, including 45.84% uniquely mapped reads and 14.51% mapped to multiple genomic locations. The resulting alignment files were processed using SAMtools v1.21 (RRID:SCR_002105) to extract strand-associated alignments, which were used as transcriptomic evidence for gene model prediction. Protein evidence was derived from curated Metazoa proteomes retrieved from OrthoDB v11 (https://bioinf.uni-greifswald.de/bioinf/partitioned_odb11/) and incorporated into the annotation workflow in accordance with the recommendations of the BRAKER pipeline [17].

As part of the funannotate v1.8.5 (RRID:SCR_023039) annotation workflow, five ab initio gene predictors—AUGUSTUS v3.1.0 (RRID:SCR_008417), SNAP v2006-07-28 (RRID:SCR_007936), glimmerHMM v3.0.1 (RRID:SCR_002654), CodingQuarry v2.0-11, and GeneMark-ES/ET v4.72_lic—were run independently [42–47]. Their predictions were subsequently integrated into a consensus gene set using EVidenceModeler (EVM) v2.1.0 (RRID:SCR_014659) [48], which combines ab initio predictions with transcriptomic and protein evidence to reconcile competing gene models and produce consensus gene annotations. Strand-specific RNA-seq alignments provided transcriptional support for exon–intron boundaries and splice junctions, whereas the curated Metazoa protein set supplied complementary homology-based evidence for predicted gene models.

Transfer RNA (tRNA) genes were predicted using tRNAscan-SE v2.0.12 (RRID:SCR_008637) [42]. To distinguish potentially functional tRNA genes from pseudogenes and other tRNA-like sequences, the initial set of 13,101 candidate predictions was subjected to the EukHighConfidenceFilter procedure implemented in tRNAscan-SE. This post-processing step evaluates candidate predictions using domain-specific covariance model scores, secondary-structure scores, and isotype-specific model scores, followed by additional filtering based on isotype and anticodon consistency. Using the default filtering thresholds, 1,445 predictions (11.03% of the initial candidates) were classified as high-confidence tRNAs. These high-confidence tRNA predictions were then integrated with the selected protein-coding gene annotation in GFF3 format using AGAT v1.4.1 (RRID:SCR_027223) [49] to generate a unified structural annotation. Following downstream annotation curation and integration into the final structural annotation, 1,383 tRNAs were retained in the final GFF3 annotation. The reduction reflects the removal of overlapping or redundant tRNA predictions during curation and subsequent NCBI indexing. The resulting GFF3 file was then screened for overlapping gene models using AGAT v1.4.1 (RRID:SCR_027223) and subsequently validated with GenomeTools v1.6.5 (RRID:SCR_016120) [41]. An additional BUSCO analysis was run on the predicted proteome of this finalised structural annotation file. Gffread v0.12.7 (RRID:SCR_018965) [50] was employed to extract the predicted protein sequences from the GFF3 annotation for functional annotation.

All predicted proteins were functionally annotated using multiple annotation sources including InterProScan v5.59-91.0 (RRID:SCR_005829) [51], EggNOG-mapper v2.1.12 (RRID:SCR_021165) [52,53], and Phobius v1.01 (RRID:SCR_015643) [54]. The respective annotation files and predicted genes were then fed into funannotate annotate to integrate the functional annotations into the final gff3 file.

## Availability of source code and requirements

Project name: *Cyphastrea microphthalma* (Cmic) isolate UAE_SA_Cmic_10 reference genome assembly and annotation.

Project home page: https://github.com/SequAna-Ukon/Cyphastrea_microphthalma_genome

Operating system(s): Linux

Programming language: Bash

Other requirements: The analysis pipelines require the bioinformatic tools and software dependencies listed in Table 4. Detailed installation instructions, pipeline descriptions, and commands are provided in the GitHub repository.

**Table 4.** Software tools and databases: versions and sources. In order of appearance in main text; databases marked with asterisks.

| Software tool | Version | Source |
|---|---|---|
| Smudgeplot | 0.4.0 | https://github.com/KamilSJaron/smudgeplot |
| GenomeScope2.0 | 2.0 | https://github.com/tbenavi1/genomescope2.0 |
| GeneForge | 1.0 | https://github.com/SequAna-Ukon/GeneForge |
| BRAKER | 3.0.8 | https://github.com/Gaius-Augustus/BRAKER |
| funannotate | 1.8.5 | https://github.com/nextgenusfs/funannotate |
| BUSCO* | 6.0.0 | https://busco.ezlab.org/busco_userguide.html |
| MinKNOW | 23.07.12 | https://nanoporetech.com/document/experiment-companion-minknow |
| Dorado | 0.7.4 | https://github.com/nanoporetech/dorado |
| FastK | 1.1 | https://github.com/thegenemyers/FASTK |
| porechop | 0.2.4 | https://github.com/rrwick/Porechop |
| NanoPlot | 1.44.1 | https://github.com/wdecoster/NanoPlot |
| Flye | 2.9.5 | https://github.com/mikolmogorov/Flye |
| BlobToolKit | 4.3.0 | https://github.com/genomehubs/blobtoolkit |
| BLAST | 2.16.0 | https://ftp.ncbi.nlm.nih.gov/blast/executables/LATEST/ |
| DIAMOND | 2.1.10 | https://github.com/bbuchfink/diamond |
| Minimap2 | 2.28-r1209 | https://github.com/lh3/minimap2 |
| Medaka | 1.12.0 | https://github.com/nanoporetech/medaka |
| Pilon | 1.24 | https://github.com/broadinstitute/pilon |
| Redundans | 2.0.1 | https://github.com/Gabaldonlab/redundans |
| GetOrganelle | 1.7.7.1 | https://github.com/Kinggerm/GetOrganelle |
| MITOS2 | 2.1.9 | https://gitlab.com/Bernt/MITOS/ |
| RepeatModeler | 2.0 | https://github.com/Dfam-consortium/RepeatModeler |
| EDTA | 2.2.2 | https://github.com/oushujun/EDTA |
| RepeatMasker | 4.1.6 | https://github.com/Dfam-consortium/RepeatMasker |
| FastQC | 0.12.1 | https://github.com/s-andrews/FastQC |
| Trimmomatic | 0.39 | https://github.com/usadellab/Trimmomatic |
| STAR | 2.7.11b | https://github.com/alexdobin/STAR |
| SAMtools | 1.21 | https://github.com/samtools/samtools |
| AUGUSTUS | 3.1.0 | https://github.com/gaius-augustus/augustus |
| SNAP | 2006-07-28 | https://github.com/KorfLab/SNAP |
| glimmerHMM | 3.0.1 | https://github.com/kblin/glimmerhmm |
| CodingQuarry | 2.0-11 | https://github.com/UBTEC/C3Q |
| GeneMark-ES/ET | 4.72_lic | https://github.com/gatech-genemark/GeneMark-EP-plus |
| EVidenceModeler | 2.1.0 | https://github.com/EVidenceModeler/EVidenceModeler |
| tRNAscan-SE | 2.0.12 | https://github.com/UCSC-LoweLab/tRNAscan-SE |
| AGAT | 1.4.1 | https://agat.readthedocs.io/en/latest/index.html |
| GenomeTools | 1.6.5 | https://github.com/genometools/genometools |
| gffread | 0.12.7 | https://github.com/gpertea/gffread |
| UniProt Reference Proteomes database |  | https://ftp.uniprot.org/pub/databases/uniprot/current_release/knowledgebase/reference_proteomes/ |
| NCBI nucleotide (nt) database |  | https://ftp.ncbi.nlm.nih.gov/blast/db/ |
| InterProScan* | 5.59-91.0 | https://github.com/ebi-pf-team/interproscan |
| OrthoDB | 11 | https://github.com/tomasbruna/orthodb-clades |
| EggNOG-mapper* | 2.1.12 | https://github.com/eggnogdb/eggnog-mapper |
| Phobius* | 1.01 | https://phobius.sbc.su.se/ |
| SignalP* | 6.0 | https://github.com/fteufel/signalp-6.0 |
| COG* | 2024 | https://www.ncbi.nlm.nih.gov/research/cog/ |
| Pfam* | 37.4 | https://pfam.xfam.org/ |
| The Gene Ontology (GO) knowledgebase* | 2025-07-22 | https://www.geneontology.org/ |
| MEROPS* | 12.5 | https://www.ebi.ac.uk/merops/ |
| the Carbohydrate-Active enZYmes Database* | 2025 | https://www.cazy.org/ |
| Kyoto Encyclopedia of Genes and Genomes (KEGG)* | 2025 | https://www.genome.jp/kegg/ |

License: MIT License

## Data Availability

Raw sequencing data (gDNA and RNA-Seq), the genome (and mitogenome) assembly and annotation (GenBank accession numbers: GCA_059441185.1, https://www.ncbi.nlm.nih.gov/datasets/genome/GCA_059441185.1; CM182515.1, https://www.ncbi.nlm.nih.gov/nuccore/CM182515.1) are deposited at NCBI under the BioProject PRJNA1190223 (gDNA; https://www.ncbi.nlm.nih.gov/bioproject/PRJNA1190223) and PRJNA1354406 (RNA-Seq; https://www.ncbi.nlm.nih.gov/bioproject/1354406) within the Umbrella BioProject PRJNA749006 (https://www.ncbi.nlm.nih.gov/bioproject/749006). All bioinformatic tools used to assemble and annotate the *Cyphastrea microphthalma* genome can be found in Table 4. The GitHub repository is published on Zenodo under https://zenodo.org/records/22676555 [55].

## Ethics Approval & Consent for Publication

Not applicable.

## Acknowledgements

We thank Dain McParland for field support, and we thank Luigi Colin for technical support.

## Author contributions

CRV conceived and conceptualized the study. RA, AF, GP, HM, CRV collected the samples. JAB provided field support. RA and AF processed the samples. AS assembled and annotated the genome. AS visualized the results. RA, AS, and CRV interpreted all analyses and results. RA wrote the original draft. All authors edited and approved the final manuscript.

